# Hypophosphite is a naturally-occurring selective inhibitor of syntrophic methanogenesis

**DOI:** 10.64898/2026.03.11.711166

**Authors:** Ruiwen Hu, Matt E. Weaver, Leslie A. Day, John Mark Marquez, Heidi S. Aronson, David A.O. Meier, Pedro Romero, Farid Halim, Alesha D. Maxwell, Kyle N. Costa, Adam M. Deutschbauer, Morgan N. Price, Matthias Hess, Koushik Singha Roy, Ando Radanielson, John D. Coates, Nicolas Tsesmetzis, Hans K. Carlson

## Abstract

Microbial methanogenesis is a major contributor to global warming and methane fluxes represent a loss of energy and electrons from industrial ecosystems. The chemical space of methane control strategies is still under-explored. Most known methanogenesis inhibitors target methanogenic archaeal enzymes. However, interference with syntrophic electron exchange in methanogenic systems presents an additional target for methane control. Here we show that hypophosphite (H_2_PO_2_^-^), an inorganic formate analog, is a potent and selective inhibitor of syntrophic methanogenesis versus primary fermentation in rice field sediments and cattle rumens. Hypophosphite is also generally recognized as safe and relatively non-toxic to plants and animals. Genetic screens and physiological assays in the model methanogen *Methanococcus maripaludis* S2 implicate formate metabolism as the target of hypophosphite inhibition. Currently, there is no known biological pathway for anaerobic hypophosphite oxidation and hypophosphite is stable in anoxic sediments for weeks to months. Given its widespread natural occurrence, we propose that hypophosphite may modulate carbon cycling in natural environments. Taken together, our results suggest that hypophosphite could be used as a safe, inexpensive, strategy for methane control in syntrophic methanogenic ecosystems.

## Introduction

Microbial methanogenesis is responsible for over half of anthropogenic methane emissions. Much of this methane is derived from managed ecosystems where interventions are possible to minimize this flux^1^. Re-directing fermentative anaerobic systems towards increased accumulation of intermediates is recognized as an important approach to waste valorization and enhancing capture of energy in anaerobic digesters^2–4^ and ruminant agriculture^5^.

Most known methanogenesis inhibitors target the conserved archaeal enzyme responsible for methane synthesis, methyl coenzyme A reductase (McrA)^6^. McrA inhibitors include bromoform (CH_3_Br, the active methanogenesis inhibitor in red seaweeds)^7^, 2-bromoethanesulfonic acid (BES)^8^, and 3-nitroxypropanol (3-NOP)^9^. These compounds are selective inhibitors of methanogenesis, but issues with stability, cost and negative impacts on environmental systems limit widespread use. As such, there is an unmet need for cheap, non-toxic, potent and selective inhibitors of methanogenesis.

A large share of global microbial methane is produced as a consequence of syntrophic methanogenesis^1^. Thus, chemical inhibitors of syntrophic bacteria or interference with acetate, hydrogen or formate exchange represent viable targets for methane flux control. Fluoroacetate inhibits acetate metabolism in acetoclastic methanogens, but is very toxic and therefore unlikely to be selective enough to be useful in natural systems^10^. Cyanide (CN) or carbon monoxide (CO) are inhibitors of hydrogenase enzymes^11,12^ and hypophosphite (H_2_PO_2_^-^) is known to inhibit the bacterial molybdoenzyme formate dehydrogenase (FDH)^13,14^. These compounds can inhibit some syntroph-methanogen co-cultures.^15^ However, their mechanisms of inhibition are poorly understood, and it remains unclear how selectively these compounds target syntrophic methanogenesis versus primary fermentation.

Both CN and CO are generally toxic to people and animals because they bind to respiratory metalloenzymes^16,17^. These compounds are also microbially degraded^18,19^. As such, CN and CO are unlikely to be useful for methane control in ruminants, rice fields or reactors. In contrast, hypophosphite is largely non-toxic to humans and animals with an LD_50_ in rats of 1.6 g/kg^20^. Over the past 150 years, hypophosphite has been tested as a treatment for tuberculosis, a food preservative, and as a phosphorus source for animals^20^. While ineffective as a human therapeutic or animal phosphorus source, hypophosphite is used occasionally in food preservation and as a soluble counter-ion in bovine calcium supplements. As such, the United States Food and Drug Administration designates hypophosphite as Generally Recognized as Safe (GRAS) for use as a nutrient or dietary supplement^20^.

Given its structural similarity to formate, hypophosphite has been used as an active-site probe to study formate metabolic enzymes in fermentative bacteria. Enzyme studies show that hypophosphite is a formate-competitive, irreversible inhibitor of *Escherichia coli* formate dehydrogenase (FDH)^21^ and pyruvate formate lyase (PFL)^22^. The precise mechanism of irreversible FDH inhibition is not well characterized, but, in the absence of formate, sub-micromolar (µM) hypophosphite concentrations are sufficient to completely inactivate all FDH in *E. coli*^13^. PFL is inactivated by hypophosphite through the formation of a covalent C-P adduct at low µM concentrations^22^.

Despite the low toxicity of hypophosphite to animals and potential to interfere with microbial formate metabolism, it is unknown if hypophosphite is selectively inhibitory of syntrophs and methanogens versus primary fermentative bacteria. All studies that have used hypophosphite to study formate metabolism in methanogens and primary fermenters have used millimolar concentrations of hypophosphite^15,23^ which is well above the thermodynamically favorable formate concentration thresholds for syntrophy. The one study that quantified hypophosphite inhibitory potency against a syntroph measured a low micromolar IC_50_ under non-syntrophic, crotonate disproportionating conditions.^14^ Studies that evaluated hypophosphite inhibition of formatotrophic methanogens report variable sensitivity and only at high millimolar concentrations^24,25^. Overall, based on evidence in the literature, it is unclear whether or not hypophosphite can effectively and selectively inhibit methane production in complex, natural methanogenic microbiomes at low micromolar (µM) concentrations.

Recent work suggests hypophosphite is widely produced as part of a natural phosphorus redox cycle.^26^ Hypophosphite is consistently measured in anaerobic wetland porewater at concentrations in the high nanomolar to low µM range^26^. Strikingly, hypophosphite concentrations can exceed 350 µM in the termite gut^27^. Some work supports a biological mechanism of phosphorus reduction in environmental systems but the enzymes involved remain unknown^26,28–30^. Recent work from our group demonstrates that phosphite can be anaerobically oxidized coupled to methanogenesis via a lithosyntrophic association between phosphite-oxidizing bacteria and methanogens^31^. Although natural hypophosphite concentrations measured in wetlands and termite guts exceed those necessary to inhibit the *E. coli* FDH^13^ it is not known if hypophosphite can influence anaerobic carbon cycling in these systems or if it shows selectivity against formate metabolism in complex systems.

To bridge these knowledge gaps, we tested whether hypophosphite can serve as a selective methane-inhibition strategy in anaerobic ecosystems. More specifically, we measured the influence of hypophosphite on a genetically-tractable methanogen, on methanogenic microbiomes from rice field sediment microcosms, and on methane fluxes from rumen fluid incubations and potted rice plants. Our results implicate syntrophic formate exchange as a target of hypophosphite and we find that hypophosphite is selectively inhibitory of syntrophic methanogenesis versus primary fermentation. We also show that hypophosphite is stable in anoxic systems and can inhibit methane fluxes from potted rice plants and rumen fluid of dairy cows. Finally, comparison of the inhibitory potency of hypophosphite with measured natural concentrations in anoxic ecosystems suggests hypophosphite could be a natural control on anaerobic carbon cycling pathways and methane fluxes. Taken together, our results have implications for methane control in agricultural ecosystems and point to important future directions of inquiry for improving our understanding of the interactions between carbon and phosphorus biogeochemical cycles.

## Results

### Hypophosphite is a potent inhibitor of a model formatotrophic/hydrogenotrophic methanogen

In previous work, high millimolar hypophosphite was found to inhibit formatotrophic methanogens^24,25^. However, in these experiments, formate concentrations were also in the high millimolar range and subsequent work with *Methanococcus maripaludis* S2 (S2) has demonstrated that the formate metabolic pathway, including the formate transporters and FDH is regulated by the availability of formate in the culture media.^32,33^ We thus hypothesized that it would be possible to measure a lower half-maximal inhibitory potency (IC_50_) for hypophosphite by pre-culturing cells on formate before transferring to fresh media without formate. To test this hypothesis, we pre-cultured S2 in a chemically defined media with either hydrogen or formate as the sole electron donor and transferred these cultures into fresh media with varying concentrations of hypophosphite **(Materials and Methods, Supplementary Table S1)**. Formate pre-grown S2 transferred to fresh media with 120 mM formate as the electron donor is very insensitive to hypophosphite with an IC50 >100 mM, but these same formate pre-grown cells are very sensitive to hypophosphite when transferred to cultures with hydrogen as the sole electron donor **(Figure 1A)**. Cells pre-grown on hydrogen and transferred to hydrogen have a hypophosphite IC_50_ ∼10mM. Taken together, these results support a model in which hypophosphite is a potent inhibitor of formatotrophic methanogens only when both methanogen formate transporters are expressed and formate concentrations are low. This scenario is what we would expect to find in most natural and engineered syntrophic methanogenic systems^34,35^.

**Figure 1.**
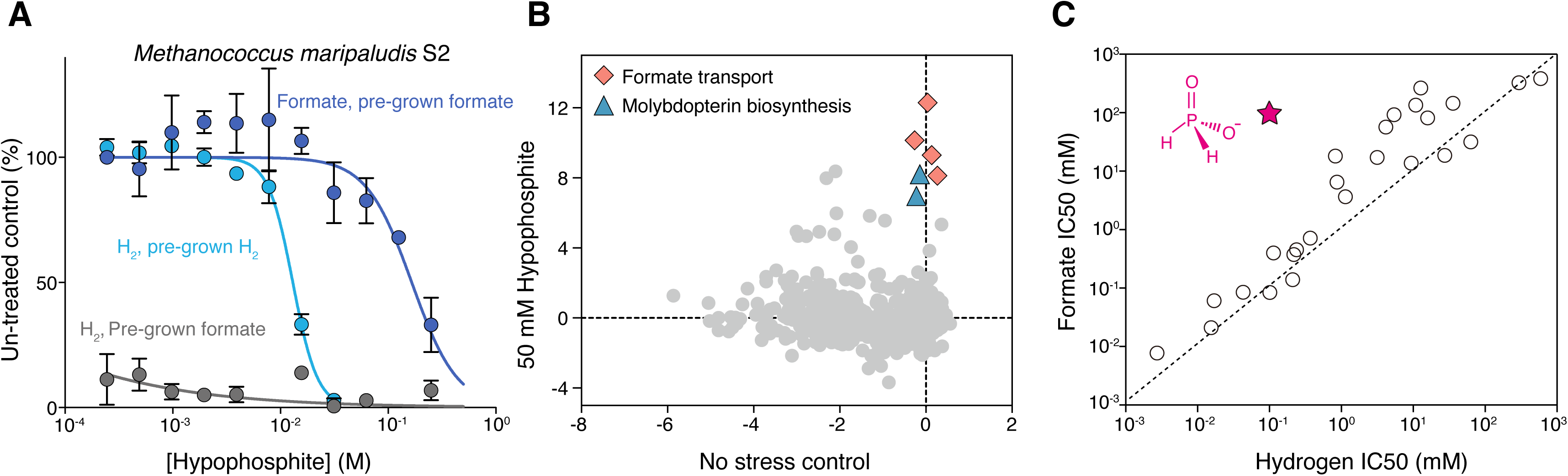
Formate metabolism and hypophosphite sensitivity in a model methanogen. **A.** Dose-response curves showing the influence of varying concentrations of hypophosphite on *Methanococcus maripaludis* S2 growth. Different cultures were pre-grown on or transferred into conditions with either hydrogen or formate as the sole electron donor. Dark blue points are cultures pre-grown on formate then transferred to formate, Light blue points are cultures pre-grown on hydrogen, then transferred to hydrogen. Dark grey points are cultures pre-grown on formate then transferred to hydrogen. Points are averages and error bars represent standard deviations of biological replicates. **B.** Genome-wide gene fitness for *Methanococcus maripaludis* S2 grown on hydrogen with no inhibitor or in the presence of 50 mM hypophosphite. Colored points indicate genes involved in molybdopterin biosynthesis, MoeA (blue) and formate transport (pink). Gene fitness is the log2 ratio of the abundance of transposon insertion strains in a gene for a given condition relative to the “time-zero” strain abundance (Materials and Methods). **C.** Half-maximal inhibitory potencies (IC_50_) for a set of compounds against *Methanococcus maripaludis* S2 grown with either formate or hydrogen as the sole electron donor. The red star represents sodium hypophosphite. The dashed line represents the same IC_50_ in formate and hydrogen grown cells.

### Methanogen formate metabolic enzymes are implicated in the mechanism of hypophosphite inhibition

To identify genes important and detrimental for *Methanococcus maripaludis* S2 growth in the presence of hypophosphite we grew a pooled bar-coded transposon mutant library^36^ **(Materials and Methods)** of S2 with hydrogen as the sole electron donor in the presence of inhibitory concentrations of hypophosphite. We found that mutants with transposon insertions in genes involved in formate transport and molybdopterin biosynthesis grew much better than the average gene indicating that these genes are strongly detrimental in the presence of hypophosphite **(Figure 1B).**

Formate transporters are involved in the uptake of formate from the extracellular environment to the cytoplasm and molybdopterin is a co-factor in the major formate metabolic enzymes, formylmethanofuran dehydrogenase (Fmd, MMP_RS02690-MMP_RS02710; Fwd, MMP_RS06425-MMP_RS06400) and formate dehydrogenase enzymes (FDH1, MMP_RS00795-MMP_RS00800; FDH2, MMP_RS06680-MMP_RS06685). We did not observe strong differential fitness during growth on hypophosphite for any of these major formate metabolic enzymes, but this may be because the multiple paralogs of these formate enzymes are potentially redundant. We were able to confirm the sensitivity of individual strains lacking two MoeA genes (MoeA, MMP_RS08320; MoeA3, MMP_RS02885) **(Figure S1).**

There are several possible explanations for the fitness advantage of molybopterin biosynthesis enzymes in resistance to hypophosphite. Hypophosphite may influence regulation of molybdopterin biosynthesis and/or directly react with molydopterin co-factor or molybdopterin enzymes. If hypophosphite is oxidized by oxidized molybdopterin the product would likely be phosphite or phosphate whereas if hypophosphite is reduced the product would be phosphine. Conceivably either of these reactions could be detrimental for fitness as hypophosphite oxidation would disrupt electron transfer, while hypophosphite reduction would produce the toxic gas phosphine which could react with metalloenzymes in the cell. While the precise chemical mechanism of potent irreversible inhibition of FDH by hypophosphite is unknown^21^ it likely involves a redox reaction. Regardless, our results implicate methanogen formate metabolism in the mechanism of hypophosphite inhibition.

### Hypophosphite is a uniquely selective inhibitor of formatotrophic methanogenesis

To determine if other compounds are similar to hypophosphite in their selective inhibition of formate metabolism in *Methanococcus maripaludis* S2 we measured the inhibitory potency of a panel of inorganic ions against S2 cultured under either hydrogenotrophic or formatotrophic growth conditions. For these assays, S2 was pre-grown on formate, then transferred into either formate or hydrogen media and mixed with serially-diluted compounds in 384-well microplates. By measuring growth relative to uninhibited control cultures, we determined the IC_50_s of diverse compounds.

Most compounds have similar IC_50_s between hydrogen and formate growth conditions, but hypophosphite is much less inhibitory for formate grown cells. (**Figure 1C** and **Supplementary Table S2)**. This result supports the view that hypophosphite is a uniquely selective inhibitor of methanogen formate metabolism.

### Hypophosphite is a selective inhibitor of methanogenesis in fermentative, anaerobic microcosms from rice field sediment

Based on the potent inhibition of *Methanococcus*, we next examined whether hypophosphite can selectively suppress methane production in a complex microbiome. We amended anoxic chemically defined microbial growth media with rice field sediment and a complex fermentable electron donor/carbon source in the form of 1 g/L yeast extract **(Materials and Methods, Supplementary Table S3)**. We passaged these microcosms three times to dilute away sediment until we could measure growth by monitoring optical density (OD600) and then transferred these enriched microcosms into fresh cultures containing varying concentrations of sodium hypophosphite. We monitored methane, growth and methanogen abundance using 16S rDNA amplicons at the early stationary phase of methane production in control and treated cultures after 3 weeks. We found that methane and the relative abundance of methanogens was inhibited with an IC_50_ of ∼50 µM hypophosphite, but optical density, indicative of fermentative growth, was not inhibited by at least 10 mM sodium hypophosphite **(Figure 2A)**. This result shows that hypophosphite can selectively inhibit methanogenesis versus primary fermentation in a complex fermentative anaerobic microbiome.

**Figure 2.**
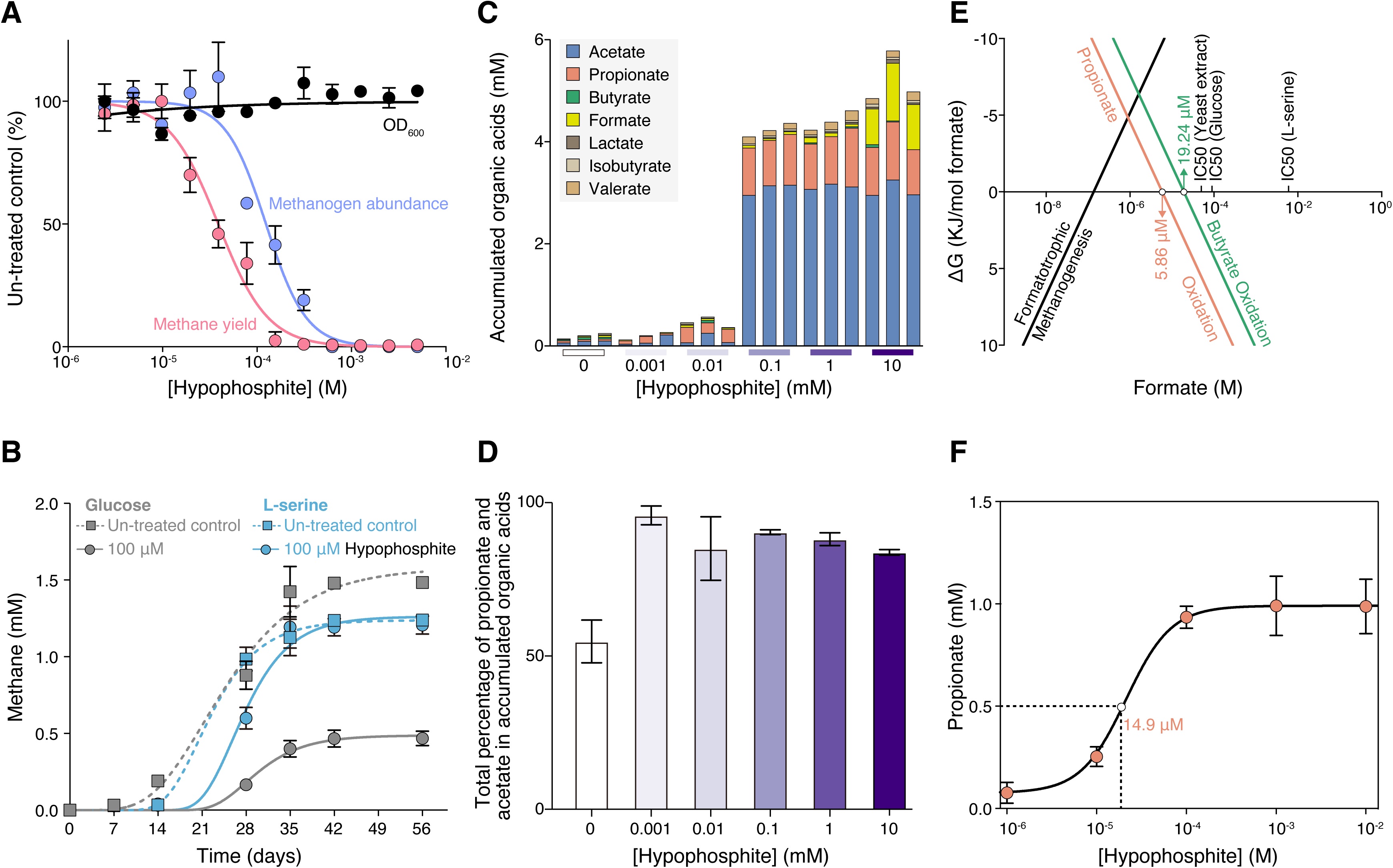
Inhibition of methanogenesis by hypophosphite in syntrophic methanogenic microcosms from rice fields. **A.** Dose-response curves showing the influence of varying concentrations of hypophosphite on methane production, growth yield (OD 600) and the relative abundance of methanogenic archaea (via 16S rDNA amplicon sequencing) in fermentative methanogenic rice field microcosms with yeast extract as the sole electron donor. Points are averages and error bars represent standard deviations of biological replicates. **B.** Hypophosphite inhibition of methane production in replicate microcosms with D-glucose (light grey) or L-serine (sky-blue) as the sole electron donor. **C.** Organic acids after 35 days in D-glucose microcosms with varying concentrations of hypophosphite Points are averages and error bars represent standard deviations of biological replicates. **D.** Propionate and acetate as a proportion of total organic acid electrons in D-glucose microcosms. **E.** Overall Gibbs energy (ΔG_r_, kJ/mol formate) of formatotrophic methanogenesis or syntrophic butyrate or propionate oxidation (37 °C) as function of formate concentrations compared with IC50s against methane yields in rice field microcosms. **F.** Propionate concentrations measured in methanogenic D-glucose microcosms as a function of hypophosphite concentrations compared with the free energy of syntrophic propionate oxidation as a function of formate concentrations. Half-maximal concentrations of propionate accumulation are shown with dotted lines. Points are averages and error bars represent standard deviations of biological replicates.

### Hypophosphite is more inhibitory of methanogenic systems that rely more heavily on syntrophic electron exchange

Syntrophic organic acid oxidation to formate becomes thermodynamically unfavorable above concentrations of ∼10 µM formate.^34^ Thus, we hypothesized that interference with formate electron exchange is an important mechanism of hypophosphite inhibition. We also hypothesized that the degree to which a methanogenic system relies on syntrophic electron exchange influences hypophosphite sensitivity.

To evaluate the importance of syntrophy in controlling hypophosphite inhibitory potency, we prepared rice field sediment microcosms with electron donors that vary in nominal oxidation state of organic carbon (NOSC). We previously demonstrated that fermentative methanogenic degradation of more reduced organic carbon (negative or zero NOSC) relies more heavily on syntrophic methanogenesis compared to degradation of more oxidized organic carbon^37^. Consistent with our hypothesis, we found that hypophosphite is a more potent inhibitor of methanogenesis in microcosms amended with D-glucose (NOSC=0) compared to microcosms amended with L-serine (NOSC = +1). After 56 hours of incubation with 100 µM hypophosphite, methane production in D-glucose-amended microcosms was reduced to ∼25% of the un-treated controls, whereas methane production in L-serine-amended microcosms was less affected **(Figure 2B)**. Notably, D-glucose is a dominant electron donor in rice field sediments^38^ and ruminants and syntrophy is important in both of these systems.

### Hypophosphite likely competes with formate exchange in syntrophic methanogenesis

To further evaluate the competitive interference of hypophosphite with syntrophic formatotrophic methanogenesis, we measured organic acid concentrations after 28 days in D-glucose and L-serine cultures in the presence of varying concentrations of hypophosphite. We observed accumulation of acetate, propionate, butyrate and other minor organic acids in D-glucose cultures amended with 100 µM or more hypophosphite (**Figure 2C, Supplemental Table S4).** We also observed organic acid accumulation at 10 mM hypophosphite in L-serine cultures (**Supplemental Table S5)**. These results are further evidence of inhibition of syntrophic methanogenesis while primary fermentation proceeds. Additionally, we found that propionate and acetate accumulate in D-glucose cultures relative to other organic acids at concentrations of 1 µM hypophosphite and above (**Figure 2D)**. In diverse terrestrial wetlands, hypophosphite concentrations can reach 2 µM^26^, and, therefore, our observation suggests that naturally produced hypophosphite may influence carbon flow in methanogenic microbiomes.

We next compared measured hypophosphite IC_50_ values against the thermodynamically favorable formate concentration thresholds for syntrophic butyrate and propionate oxidation. We found that IC_50_s for the D-glucose and yeast extract cultures, which rely more heavily on syntrophic electron exchange^37^, are close to the thermodynamic formate concentration thresholds (**Figure 2E)**. Direct quantification of formate at low micromolar concentrations is technically challenging. We instead examined propionate accumulation in D-glucose cultures amended with varying concentrations of hypophosphite (1 µM–10 mM). We found that the concentrations of hypophosphite equal to or above 10 µM (EC_50_ = 14.9 µM) were sufficient to drive propionate accumulation **(Figure 2F)**. This result is consistent with a model where hypophosphite interferes with formate exchange in syntrophic methanogenesis and thereby stalls syntrophic propionate oxidation.

### Hypophosphite is not a potent inhibitor of other respiratory metabolisms or methane oxidation

To further assess the selectivity of hypophosphite against syntrophic methanogenesis versus other metabolisms, we prepared rice field sediment microcosms with 1 g/L yeast extract as the sole electron donor, and added sulfate or nitrate as terminal electron acceptors. Again, after passaging these cultures multiple times to remove residual sediment that interferes with measurements of optical density, we measured growth by optical density and accumulation of nitrite or sulfide using colorimetric assays. We found that neither fermentative growth nor nitrate or sulfate reduction were inhibited in these cultures up to a concentration of 10 mM sodium hypophosphite **(Supplemental Figure S2A).** We also measured the inhibitory potency against a model aerobic methanotroph, *Methylosinus trichosporium* OB3B, grown with methane as the sole electron donor, and against a model aerobic heterotroph, *Escherichia coli* K-12. We found that these organisms had IC_50_s above 10 mM hypophosphite **(Supplemental Figure S2B).** Along with the results presented in **Figure 1** and **Figure 2**, these observation further supports the selective inhibitory effect of syntrophic methanogenesis by hypophosphite relative to primary fermentation or other respiratory metabolisms and is consistent with other work showing that growth of diverse bacteria is not greatly impacted by concentrations of hypophosphite up to 10 mM^23,39^.

### Hypophosphite inhibits methane production in rumen fluid from cows

Ruminants are responsible for around 16% of global methane fluxes^40^. As such, there is widespread interest in developing strategies for methane mitigation for ruminants. We prepared anoxic rumen enrichments in serum bottles by mixing rumen fluid with a concentrated bicarbonate buffer to neutralize organic acids and amended with finely milled cattle feed as the sole electron donor and varying concentrations of hypophosphite **(Materials and Methods)**. We found that 1 mM hypophosphite inhibited ∼25% of total methane production after 24 hours of incubation at 37 °C but higher doses did not lead to significantly more inhibition (**Figure 3A and 3B)**. This is consistent with reports that formate is responsible for ∼30% of the total methane flux from the bovine rumen ^41^. We next pre-incubated the rumen fluid in the absence of hypophosphite for 7 days and upon testing hypophosphite inhibitory potency found that 0.1 mM hypophosphite was capable of inhibition of 25% of total methane after 192 hours **(Figure 3A and 3C**). This result is consistent with inhibition of formate exchange in syntrophic methanogenesis (**Figure 1** and **Figure 2**).

**Figure 3.**
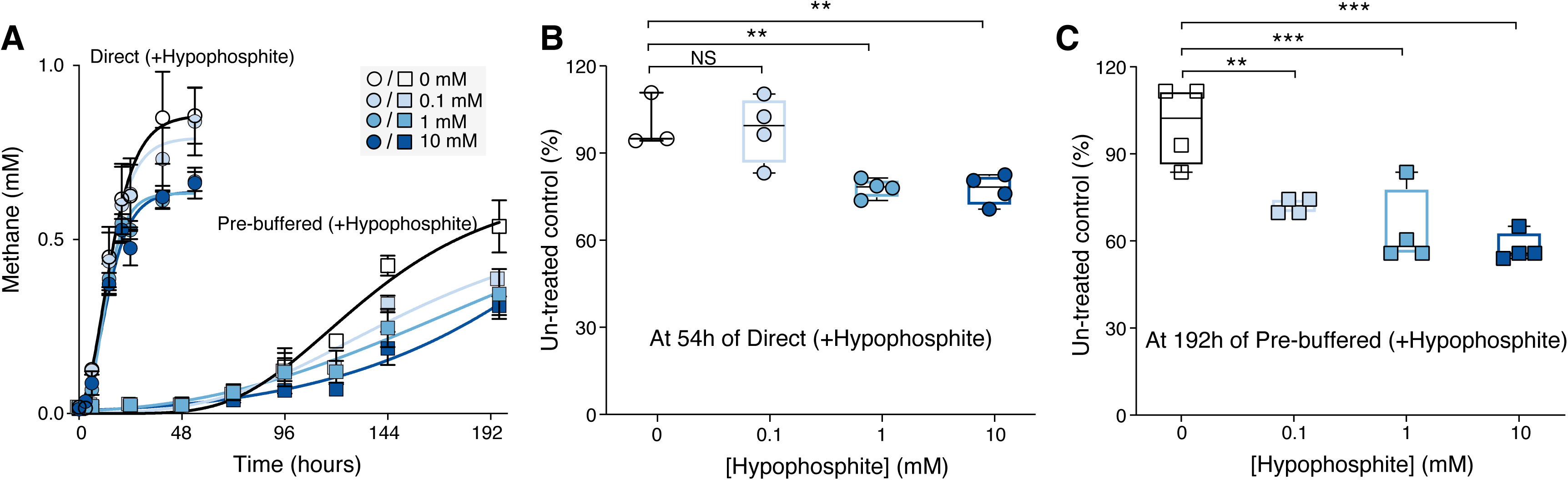
Hypophosphite inhibition of methane production in bovine rumen fluid. **A.** Methane production over time in buffered rumen fluid incubations amended with varying concentrations of hypophosphite across 0 to 10 mM. Circles are incubations where rumen fluid is added directly to a buffer containing hypophosphite. Squares are pre-buffered rumen fluid transferred into incubations with varying concentrations of hypophosphite. Points are averages and error bars represent standard deviations of biological replicates. **B and C.** Methane production relative to un-treated controls (%) for 0.1, 1 and 10 mM hypophosphite treated incubations B: Direct incubations, and C: Pre-buffered incubations. Statistics analysis of significance were performed by ANOVA, *P* <0.001 (***); *P* <0.01 (**).

### Hypophosphite inhibits methane production in potted rice plants, and remains stable in anaerobic sediments

Alongside ruminants, rice agriculture is another major global source of methane^1^. To evaluate the potential of hypophosphite to inhibit methane in rice cultivation systems we treated flooded rice pots with 0.1 mM hypophosphite and monitored methane over time. We found that methane was inhibited up to 80% by hypophosphite after 18 days (**Figure 4A)**. No impact on plant health or rice yield was observed. We also monitored the stability of hypophosphite in rice pot porewater and measured no significant loss after 18 days (**Figure 4B)**. Currently, no anaerobic microbial pathways of hypophosphite oxidation are known. However, hypophosphite is prone to abiotic oxidation in the presence of metals and oxygen and has a short half life in oxidized sediments^26^. Notably, the oxidation of hypophosphite would result in phosphate as the major end-product thereby converting this methane inhibitor into an essential nutrient for plants.

**Figure 4.**
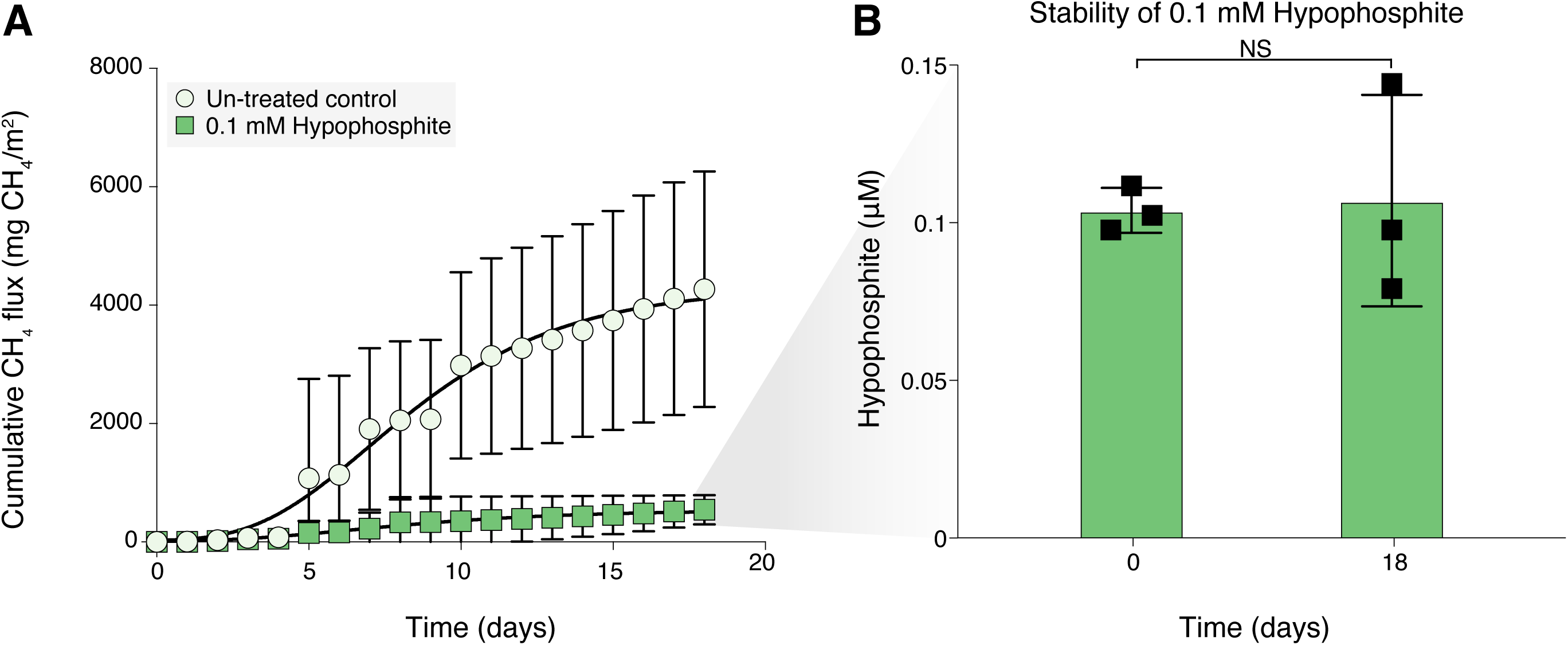
Hypophosphite inhibition of methane fluxes in potted rice plants. **A.** Cumulative methane in un-treated control rice pots (n=3) compared with 0.1 mM hypophosphite treated rice pots (n=3). Points are averages and error bars represent standard deviations of biological replicates. **B.** Hypophosphite concentrations at initial timepoint and after 18 days in rice pot porewater. Statistics analysis of significance were performed by t tests, *P* <0.05 (*).

## Discussion

Here we provide evidence that hypophosphite, an inorganic formate analog, is a potent and selective inhibitor of formate exchange during syntrophic methanogenesis. Physiological and genetic experiments implicate formate metabolic enzymes in a model methanogen as the target of hypophosphite. In complex fermentative methanogenic microbiomes, hypophosphite concentrations that exceed the thermodynamically favorable formate concentration range for syntrophy inhibit methane production and lead to accumulation of organic acids while allowing primary fermentation to proceed. Preliminary tests in complex potted rice plant mesocosms and rumen fluid from dairy cows suggest that hypophosphite could be useful for modulating methane fluxes in diverse fermentative methanogenic ecosystems where syntrophic electron exchange is important.

The apparent selectivity and potency of hypophosphite as an inhibitor of formatotrophic methanogenesis suggests that it could be a useful tool for studying the relative importance of formate as an electron carrier in diverse syntrophic methanogenic systems. While formate diffuses ∼5x more slowly than hydrogen, formate is 250-fold higher in concentration when in thermodynamic equilibrium with hydrogen^34,35^. Thus, it is thought that hydrogen is more important as an aqueous electron carrier over longer distances while formate is dominant as an electron shuttle in biofilms and methanogenic microbial aggregates such as anaerobic digester granules.^34^

Hypophosphite is an inexpensive, inorganic compound and is stable in anoxic ecosystems. In rice fields and rumen microbiomes from cows, 100 µM to 1 mM hypophosphite inhibits between 25-50% of methanogenesis. At these concentrations, hypophosphite is non-toxic to rice plants or ruminants. Because hypophosphite targets formate utilization, it is not expected to eliminate methane production entirely in systems where methanogens can use other electron carriers (e.g. hydrogen). This is consistent with our results in cow rumen fluid where ∼25-30% of methane production is suppressible and it is thought that a similar proportion of methane is derived from formate^41^. Future work will explore the potential benefits for methane inhibition from combining hypophosphite with inhibitors of hydrogenotrophic methanogenesis.

Also, future research will focus on determining if hypophosphite can function as both a methane inhibitor and phosphorus source in rice fields and as a methane control strategy in ruminant agriculture. Evaluating hypophosphite in other industrial ecosystems where methane control is important such as anaerobic digesters, manure lagoons, or subsurface hydrogen gas storage are also future areas of investigation. In forthcoming work, we have evidence that hypophosphite is effective in methane control in lab-scale anaerobic digester systems (Matt Weaver *et al.*, Unpublished results).

Additionally, our results suggest that naturally-produced hypophosphite could modulate anaerobic carbon cycling and methane fluxes in anoxic systems such as wetlands or the termite gut. Phosphorus undergoes a natural redox cycle.^26^ A survey of anoxic aquatic sediments found that reduced phosphorus represented up to a third of total phosphorus^26^, with hypophosphite concentrations up to ∼2 µM. In termite guts, hypophosphite was measured at >300 µM^27^. While the sources of reduced phosphorus in these environments are unknown, it is possible that hypophosphite could be produced through biological phosphorus reduction^26,28–30,42^.

We compared the measured inhibitory potency of hypophosphite against diverse formate metabolic enzymes and microbial processes with natural concentrations in different ecosystems **(Figure 5A)**. It is clear from this analysis that hypophosphite concentrations in aquatic sediments are sufficient to inhibit microbial formate dehydrogenase enzymes and cause a shift in syntrophic organic acid composition (as shown in **Figure 1 and Figure 2)**. In termite guts, hypophosphite concentrations exceed the inhibitory concentrations required to inhibit syntrophic methanogenesis in our microcosms from rice fields and cow rumens. Termite guts, unlike the rumen or most other metazoan guts, have a dominance of homoacetogenic bacteria over methanogenic archaea and most methane is derived from hydrogen and acetate rather than formate in termite guts^43^. The basis for homoacetogenesis in termite guts has long been debated, but based on our results we hypothesize that high hypophosphite concentrations are a plausible explanation that to our knowledge has not been discussed in the literature. By annotating formate dehydrogenase and formate transporter genes across 893 archaeal methanogen genomes from the Genome Taxonomy Database (GTDB)^44^, we found that more than 30% of methanogens possess the genetic potential for formate utilization **(Supplementary Figure 3)**. This prevalence indicates that formatotrophy is widespread among methanogenic archaea and represents a phylogenetically broad potential target for hypophosphite inhibition.

**Figure 5.**
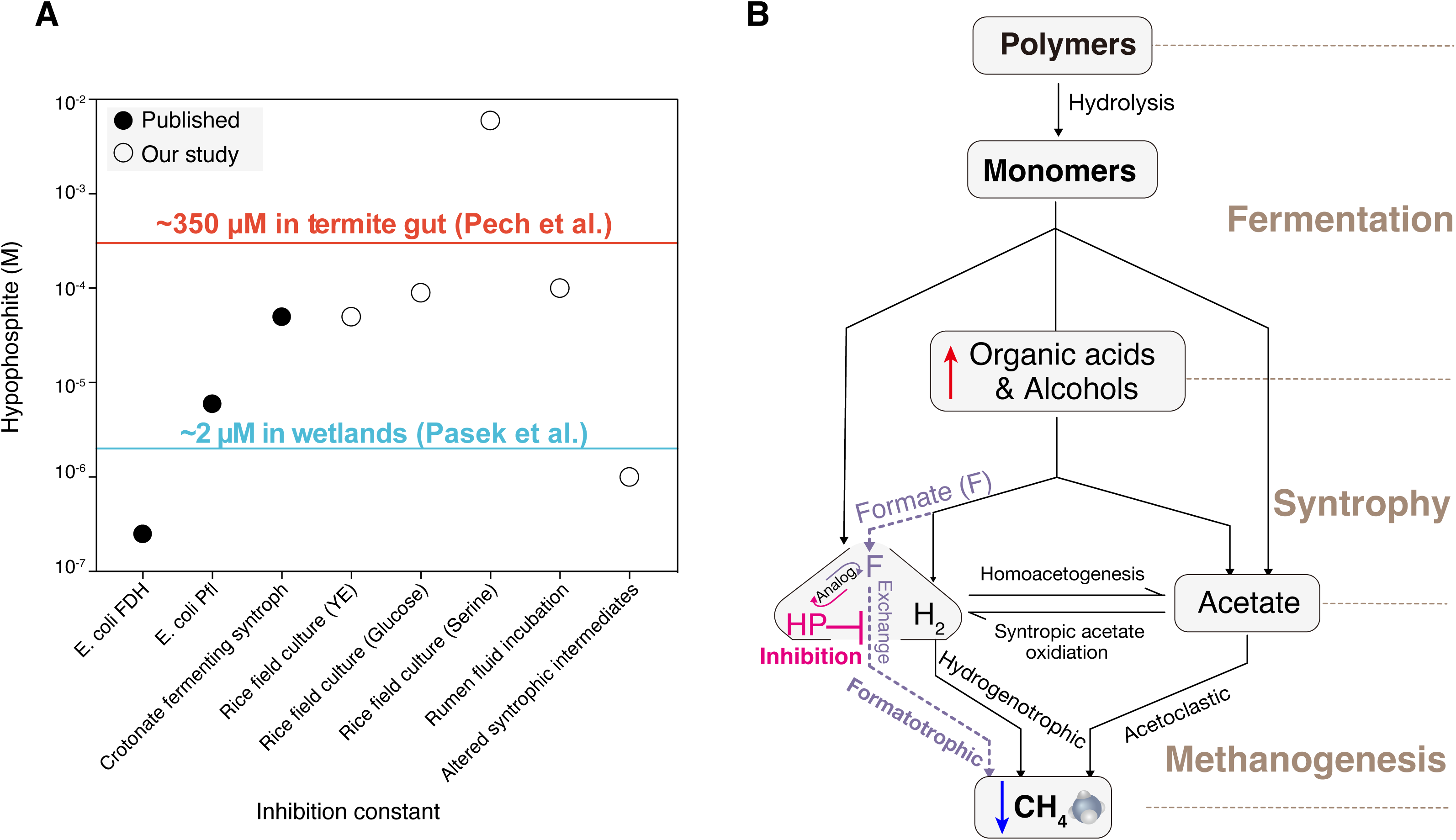
Hypophosphite as a natural, potent and selective inhibitor of formate exchange in syntrophic methanogenesis. **A.** Inhibitory potencies of hypophosphite against different microbial enzymes or microbial processes compared with environmental concentrations. **B.** Conceptual model showing likely targets of hypophosphite in complex carbon and electron flow processes in fermentative methanogenic systems.

We propose that hypophosphite selectively inhibits syntrophic methanogenesis by competing with formate for uptake by formatotrophic methanogens **(Figure 5B)** while allowing primary fermentation to proceed. While higher concentrations of hypophosphite can interfere with fermentation and respiration, the selectivity window is several orders of magnitude. Identifying and characterizing the enzymes responsible for phosphorus reduction is critical for understanding the mechanisms that regulate steady state concentrations of reduced phosphorus and its role in modulating methane flux. Understanding the interactions between the anaerobic carbon and phosphorus cycle has implications for biogeochemical modelling of these cycles.

### Environmental Implication

Most strategies for microbial methane control involve toxic, expensive and rapidly degraded organic compounds. Hypophosphite is an inorganic compound with a long half-life in the absence of oxygen. Here we provide the first direct evidence that hypophosphite can selectively interfere with syntrophic methanogenesis in rice fields and ruminants but is relatively non-toxic to primary fermentative bacteria, respiratory bacteria, plants and animals. We also propose that naturally produced hypophosphite can modulate methane fluxes in natural anaerobic sediments and termite guts, suggesting a previously overlooked interaction between the carbon and phosphorus redox cycles. This work identifies an abundant natural compound that safely suppresses methane production in anaerobic environments by targeting microbial interactions, rather than killing the microbes or blocking essential metabolic enzymes in methanogens.

## Methods

### Media and cultivation conditions

All media recipes are in **Supplementary Materials**. *Methanococcus maripaludis* S2 was cultivated in a basal media in anaerobic Balch tubes as described previously^36^ or in microplates in an anoxic pressurized incubation vessel using either formate or hydrogen as the electron donor. *E. coli* was cultured in LB broth (Difco) and in NMS media in sealed Balch tubes with a headspace of CH_4_ and air. Rice field microcosms were cultured in a chemically defined basal media as described previously^37^. Organic carbon compounds, stressors and sodium hypophosphite were added from concentrated anoxic stock solutions (Sigma-Aldrich).

### Analytical methods

Headspace methane concentrations in anaerobic Balch tubes and serum bottle cultures were measured with a Agilent gas chromatograph model 7890A (G344OA) with a Superlco SP™-2380 Fused Silica Capillary column containing a proprietary, bonded silica-based stationary phase. Methane was measured using a flame ionization detector (FID) and GC oven temperature was held at 50 °C for 60 minutes to allow for non-carrier gases to elute. The FID heater was kept at 300°C; air flow was 400mL/min; hydrogen gas fuel flow was 30mL/min; and nitrogen gas flow was 25 mL/min. (H2 and N2 are the carrier gases for FID). Methane standard gases (Scotty Analyzed Gases) were used for calibration. GC ChemStation (Agilent Technologies) computer software was used for quantification. Growth of anaerobic Balch tube and serum bottle cultures were measured using a Spectronic 20D spectrophotometer. Hypophosphite concentrations were measured using ion chromatography (Dionex).^45^ To measure the syntrophic intermediate metabolites across anaerobic Balch tubes treated with glucose, we quantified formate, acetate, lactate, propionate, butyrate, isobutyrate and valerate by high-performance liquid chromatography on a Shimadzu LC-20AD using an Aminex® HPC-87H, 300mm x 7.8mm with a 5mM sulfuric acid mobile phase at a flow rate of 0.6mL/min. The pressure and temperature were held at 720psi and 50°C. Standard curves were prepared for quantification of each organic acid.

### Dose-response assays and analysis with *Methanococcus maripaludis*

For high-throughput dose-response assays concentrated stock solutions were serially Z-diluted across three 384 well plates to attain twelve two-fold dilutions of each compound^46–48^. These arrayed dose-response plates were inoculated 1:1 with 2x concentrated basal media with *Methanococcus.* For hydrogen growth assays, plates were sealed with breathable seals and grown at 37 °C in pressure vessels with a pressure of 30 psi 80:20 H_2_:CO_2_ atmosphere. For formate growth assays, plates were sealed in permeable seals and grown in anaerobic boxes (BD). All growth vessels contained paper-towels soaked in 1 M sodium sulfide as a sulfur source and atmospheric reductant. At timepoints, vessels were opened inside anaerobic chambers (Coy) and OD600 measured with a Tecan M1000 Pro microplate reader (Tecan). Plate reads were uploaded to an in-house growth database. Early stationary phase timepoints were selected for dose-response analysis and custom scripts to fit a non-linear regression to determine half-maximal inhibitory concentrations (IC_50_) ^48^.

### Genome-wide fitness assays to identify genes important for hypophosphite resistance and sensitivity in *Methanococcus maripaludis* S2

A previously constructed bar-coded transposon mutant library of Methanococcus maripaludis S2^36^ was recovered from freezer stocks and used as previously described. All media recipes are in **Supplementary Materials**. Hypophosphite was added from concentrated aqueous stock solutions for dose-response assays. Anaerobic Balch tubes were incubated at 37 °C and growth was monitored over time by OD600. Cultures that were inhibited by hypophosphite relative to un-treated controls were allowed to grow to stationary phase before collecting pellets for measuring genome wide fitness using RB-TnSeq. We represent strain fitness as the normalized log2 ratio of counts between the treatment sample (i.e., after growth in a certain condition) and the reference “time-zero” samples^49,50^. Gene fitness is the weighted average of the strain fitness **(Supplementary Table S6)**. Gene fitness for these experiments is available on fit.genomics.lbl.gov as set15S713, set15S714, set15S711, set15S712.

### Dose-response assays and analysis rice field microcosms

Rice field microcosms were cultured in a chemically defined basal media as described previously in Balch tubes^37^. Organic carbon compounds, stressors and sodium hypophosphite were added from concentrated anoxic stock solutions (Sigma-Aldrich). Anoxic sodium hypophosphite stock solutions in water were serially-diluted in Balch tubes prior to amendment with enriched microcosms. Growth was monitored by OD600 using a Spectronic 20D spectrophotometer. Methane and organic acids were measured using methods described above in **Analytical methods.**

### Thermodynamic calculations

The Gibbs energy of reaction, ΔG_r_, was calculated for formatotrophic methanogenesis (Rxn. 1), butyrate oxidation to formate coupled to bicarbonate reduction to acetate (Rxn. 2), and propionate oxidation to formate coupled to bicarbonate reduction to acetate (Rxn. 3) at 37 °C, as described in our previous work^31^. pH was set at 7, bicarbonate concentration was set at 2.98×10^-2^ M, acetate concentration was set at 8.80×10^-5^ M, methane concentration was set at 1×10^-4^ M, butyrate concentration was set at 3.5×10^-5^ M, and propionate concentration was set at 3.8×10^-5^ M. Formate concentration ranged from 1×10^-10^ to 1 M.

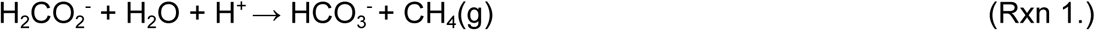

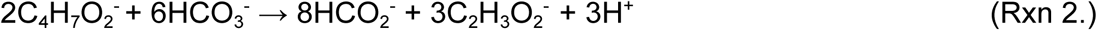

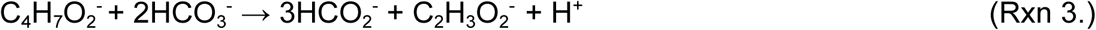

### Generation of *M. maripaludis* moeA3 deletion strain

The moeA1 deletion strain was generated in a previous publication^51^. To create the *M. maripaludis* moeA3 deletion strain, two PCR fragments were initially generated. Briefly, 1000 bp upstream and 1000 bp downstream of moeA2 gene were PCR amplified. The DNA fragments were then assembled into XbaI and NotI-digested pCRuptneo plasmid via Gibson assembly and transformed to *E. coli* DH5 via electroporation. Colonies grown on lysogeny broth solidified with 1.5% agar and ampicillin were screened by PCR to identify those carrying the correct recombinant plasmid construct. The plasmids used in this study were transformed into *M. maripaludis* S2 under anoxic conditions using a polyethylene glycol (PEG)-mediated transformation protocol. Briefly, *M. maripaludis* cultures grew to late log phase with an OD600 of ∼0.9. The cell pellet was harvested by centrifugation and washed with a transformation buffer (TB) (50 mM Tris, 350 mM sucrose, 380 mM NaCl, 1 mM MgCl2, pH 7.5). The washed cell pellet was resuspended in 0.375 ml of TB followed by addition of ∼5 µg of plasmid DNA. Next, 0.225 ml of PEG solution (40% wt/vol PEG 8000 in TB) was added to the cell mix, followed by a 1-hour non-shaking incubation at 37 °C. Transformants were washed with fresh McCas medium and pressurized to 280 kPa with an H_2_:CO_2_ (80%:20%) gas mix. After outgrowing at 37°C with agitation for at least 4 hours, transformants were sub-inoculated McCas medium containing neomycin (1 mg ml-1) to select for transformants. For counterselection of the recombinant suicide vector pCRuptneo, the transformants were plated on solid McCas medium containing 6-azauracil (0.2 g L^-1^) followed by incubation in a pressurized container with 138 kPa with a H_2_:CO_2_ (80%:20%) gas mix at 37 °C. The *M. maripaludis* colonies were screened for the deletion of moeA3 using the primers that bind ∼500 bp upstream or downstream of the moeA3 gene.

### Methane inhibition assays in rumen fluid incubations

Rumen fluid for in-vitro rumen fermentation was obtained from three rumen-fistulated non-lactating Holstein cows housed at the University of California, Davis (UCD) Dairy Teaching and Research Facility. The donor animals were fed a dry cow total mixed ration (TMR) consisting of 50% wheat, 25% alfalfa, 21.43% almond hulls, and 3.57% dry cow pellet on a dry-matter (DM) basis. Animals were fed four times a day at 5:00 am, 11:00 am, 6:00 pm, and 11:00 pm and had free access to water. Rumen fluid was collected one hour before feeding and just prior to the start of the in-vitro rumen fermentation, in accordance with the Institution of Animal Care and Use Committee (IACUC) at UCD under protocol number 22753. Approximately 500 mL of rumen fluid were collected from each cow via a perforated PVC pipe, 500 mL syringe and Tygon tubing (Saint-Gobain North America, Malvern, PA, USA), passed through a strainer, transferred into individually pre-warmed insulated thermos (Coleman Company, Chicago, IL, USA) and transported to the laboratory within 30 minutes. Equal volumes of rumen fluid from the three cows were pooled to account for biological variation, filtered through 2 layers of cheesecloth, dispensed into shipment containers, and flushed with CO_2_ for 30 minutes before sealing.

Rumen fluid was stored in insulated shipment thermos bottles until initiation of the experiment (3 hours). Rumen fluid was mixed with concentrated bicarbonate buffer^7^ and aliquoted into Balch tubes containing dry cow total mixed ration (TMR). Hypophosphite was added from concentrated aqueous stock solutions for dose-response assays. Anaerobic Balch tubes were incubated at 37 °C and methane measured over time using gas chromatography. ANOVA analysis (a significance level set at *P* < 0.05 *, *P* < 0.01 **, *P* < 0.001 ***) was used to explore the differences of hypophosphite concentrations between untreated and 0.1 mM, 1mM, 10mM hypophosphite treated Rumen fluids.

### Rice pot methane inhibition assays

Rice pot experiments were performed in the glass house at the International Rice Research Center. Two week old seedlings of IRRI 154 were transplanted into pots in a randomized complete block design. Pots were maintained in a water tank in a continuous flooded fashion. Fertilizer was added at rates of 130 kg N/ha, 50 kg P_2_O_5_/ha, and 30 kg K_2_O/ha. After 21 days of cultivation, sodium hypophosphite was applied at a rate of 0.1 mM/kg of dry soil. Methane was monitored over time with a Licor-3100 using a flux box that was placed over each pot daily between 7-11 AM for three minutes per pot. Porewater was collected for measurement of hypophosphite concentrations at 0 and 18 days of treatment and analyzed by ion chromatography as described above. Aboveground biomass (green leaves, dead leaves, stems and panicles) was measured at the end of the experiment. No significant difference in yields and health was observed between untreated and 0.1 mM hypophosphite treated rice plants. T tests (a significance level set at *P* < 0.05) were used to explore the differences of hypophosphite concentrations between untreated and 0.1 mM hypophosphite treated rice plants.

### Metabolic annotations for formate utilization across archaeal methanogens from GTDB database

To investigate the prevalence of formate utilization across diverse archaeal methanogens, we first filtered the high- or medium-quality (completeness ≥50% and contamination ≤10%) archaeal genomes from GDTB database^44^ by searching for methyl-coenzyme M reductase genes (KO ids: K00399, K00401 and K00402). A total of 893 high- or medium-quality archaeal genomes across 48 families were identified as methanogens and collected. Based on KO ids for formate dehydrogenase (K00125 and K22516) and formate transporter (K21993), we used the DRAM v1.5.0^52^ to predict the formate metabolic genes across 893 methanogenic archaea genomes **(Supplemental Table S7).** The phylogenomic tree of these methanogenic archaea was constructed by GToTree v1.8.4^53^ with default setting parameters.

## Acknowledgements

This study was financially supported by the Energy & Biosciences Institute (EBI) through the EBI-Shell program and some co-authors are members of ENIGMA (Ecosystems and Networks Integrated with Genes and Molecular Assemblies; (http://enigma.lbl.gov), a Science Focus Area Program at Lawrence Berkeley National Laboratory, US Department of Energy, Office of Science, Biological and Environmental Research Program under contract number DE-AC02-05CH11231 to Lawrence Berkeley National Laboratory.

## Author contribution

H.K.C. R.W.H., M.E.W., L.D. J.M.M. performed the laboratory work. R.W.H., H.S.A., M.N.P., L.D. and H.K.C. carried out the bioinformatics and statistical analysis. R.W.H. and H.K.C. wrote the first draft of the manuscript. All authors discussed the results. All authors read and approved the final version of the manuscript.

## Data availability

All data needed to evaluate the conclusions in the paper are present in the paper and/or the Supplementary Information. The nucleotide sequences for 16S rDNA gene amplicon sequencing and metagenomic sequencing were deposited in the SRA database under accession numbers PRJNA1188626 and PRJNA1187553, respectively. RB-TnSeq data is available on fit.genomics.lbl.gov.

## Code availability

All software packages utilized in this study are publicly accessible, and no original code is reported in this study.

