## Supplementary Materials for "Hypophosphite is a naturally-occurring selective inhibitor of syntrophic methanogenesis"

^#^corresponding author

**Supplementary Tables are in the associated .xlsx file.**

**Supplementary Table S1.** Chemically defined basal media for *Methanococcus maripaludis* S2 cultivation.

**Supplementary Table S2.** Half-maximal inhibitory potencies (IC_50_) for selected compounds against growth of *Methanococcus maripaladis* S2 grown on either H_2_ or formate after pre-culturing on formate.

**Supplementary Table S3.** Chemically defined basal media for rice field microcosm experiments.

**Supplementary Table S4.** Organic acid concentrations after 28 days in D-glucose microcosms amended with varying concentrations of sodium hypophosphite.

**Supplementary Table S5.** Organic acid concentrations after 28 days in L-serine microcosms amended with varying concentrations of sodium hypophosphite.

**Supplementary Table S6.** Genome-wide fitness data from RB-TNSeq library of *Methanococcus maripaludis* S2 for hydrogenotrophic growth in the absence and presence of 50 mM sodium hypophosphite.

**Supplementary Table S7.** Metabolic annotations for formate utilization across 893 high- or medium-quality archaeal methanogenic genomes from GTDB database.

**Supplementary Figures**.

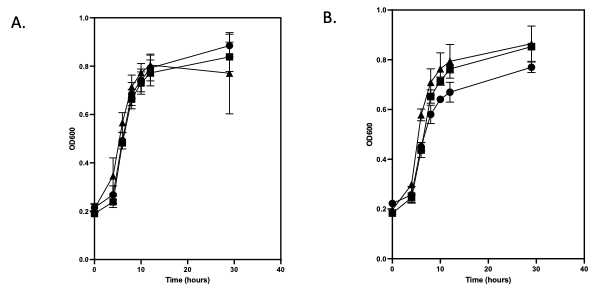

**Figure S1.** Mutants in *Methanococcus maripaludis* S2 MoeA genes have increased tolerance to hypophosphite. Growth of *Methanococcus maripladus* S2 WT (circles), MoeA (MMP_RS08320) (squares), and MoeA3 (MMP_RS02885) (triangles) in the absence **A.** or presence **B.** of 10 mM sodium hypophosphite with hydrogen as the sole electron donor.

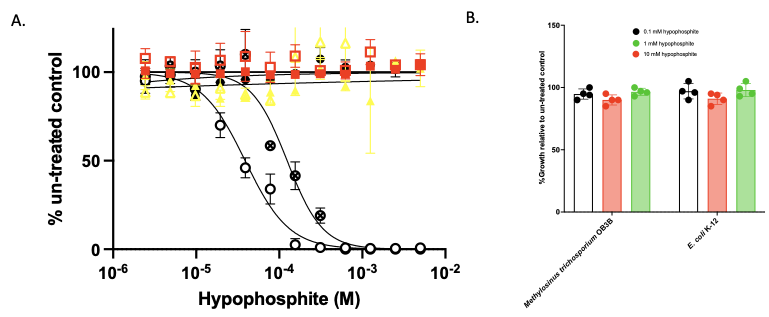

**Figure S2.** **A.** Dose-response curves showing the influence of varying concentrations of hypophosphite on methanogenic cultures (black circles) methane production (open circles), growth (OD 600) (closed circles) and the relative abundance of methanogenic archaea (x circles) as assessed using 16S rDNA amplicon sequencing in fermentative syntrophic cultures from rice fields with yeast extract as the sole electron donor. Growth (closed symbols) and respiratory activity (nitrite or sulfide production) (open symbols) for nitrate-reducing cultures (red squares) or sulfate-reducing cultures (yellow triangles). **B.** Growth relative to un-treated control cultures of *E. coli* K-12 in LB media and *M. trichosporium* OB3B in NMS media with methane as the sole electron donor.

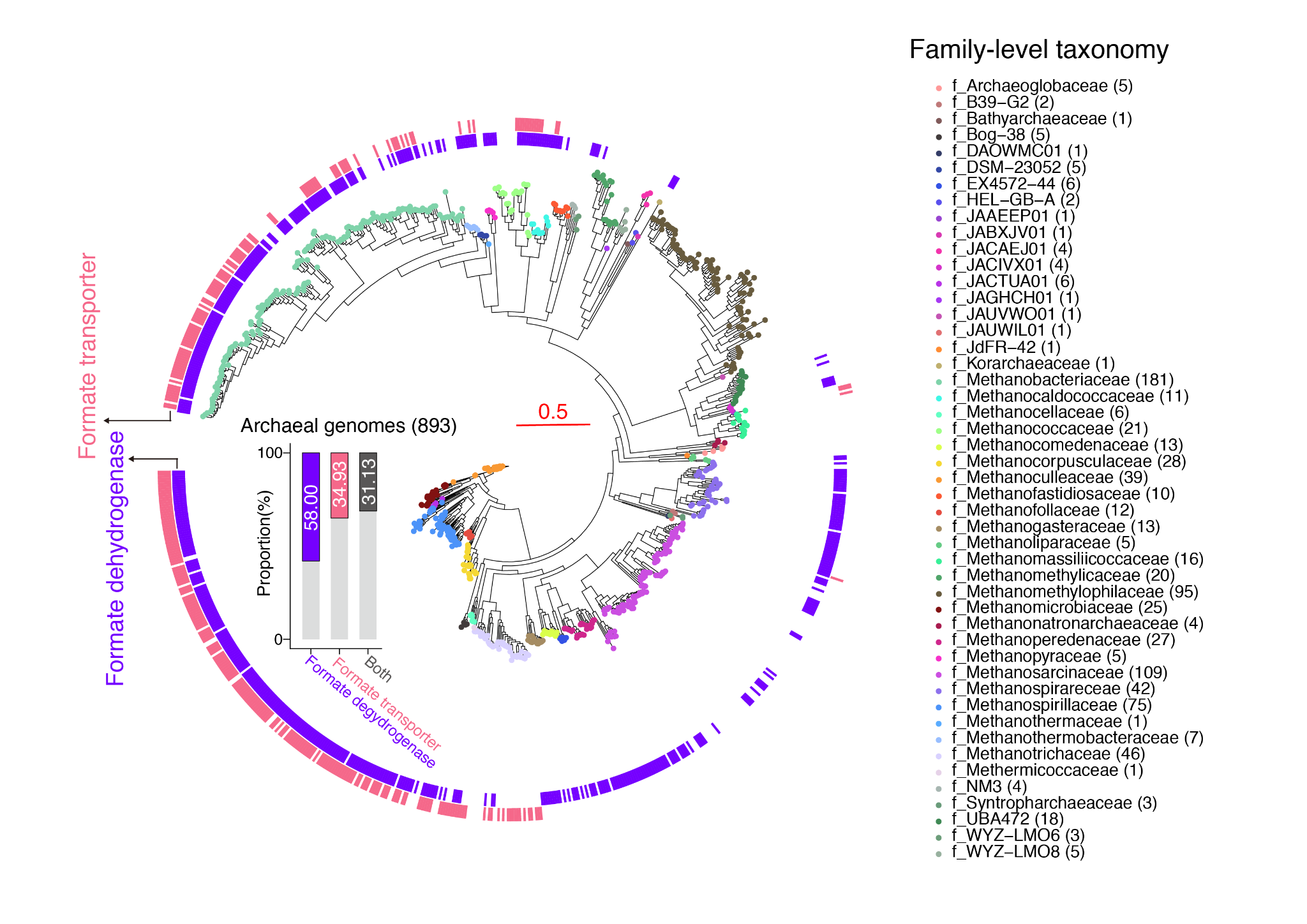

**Figure S3.** Metabolic annotations for formate utilization, including formate dehydrogenase and transporter, across 893 high- or medium-quality archaeal methanogenic genomes from GTDB database.
